# Posterior and Mid-Fusiform Contribute to Distinct Stages of Facial Expression Processing

**DOI:** 10.1101/279166

**Authors:** Yuanning Li, R. Mark Richardson, Avniel Singh Ghuman

## Abstract

Though the fusiform is well-established as a key node in the face perception network, its role in facial expression processing remains unclear, due to competing models and discrepant findings. To help resolve this debate, we recorded from 17 subjects with intracranial electrodes implanted in face sensitive patches of the fusiform. Multivariate classification analysis showed that facial expression information is represented in fusiform activity, in the same regions that represent identity, though with a smaller effect size. Examination of the spatiotemporal dynamics revealed a functional distinction between posterior and mid-fusiform expression coding, with posterior fusiform showing an early peak of facial expression sensitivity at around 180 ms after subjects viewed a face and mid-fusiform showing a later and extended peak between 230 – 460 ms. These results support the hypothesis that the fusiform plays a role in facial expression perception and highlight a qualitative functional distinction between processing in posterior and mid-fusiform, with each contributing to temporally segregated stages of expression perception.

## Introduction

Face perception, including detecting a face, recognizing face identity, assessing sex, age, emotion, attractiveness, and other characteristics associated with the face, is critical to social communication. An influential cognitive model of face processing distinguishes processes associated with recognizing the identity of a face from those associated with recognizing expression ^1^. A face sensitive region of the lateral fusiform gyrus, sometimes called the fusiform face area, is a critical node in the face processing network ^2-5^ that has been shown to be involved in identity perception ^6-10^. What role, if any, the fusiform plays in face expression processing continues to be debated, particularly given the hypothesized cognitive distinction between identity and expression perception.

Results demonstrating relative insensitivity of the fusiform to face dynamics ^11^ and reduced fusiform activity for attention to gaze direction ^12^ led to a model that proposed that this area was involved strictly in identity perception and not expression processing ^2^. This model provided neuroscientific grounding for the earlier cognitive model that hypothesized a strong division between identity and expression perception ^1^. However, some recent imaging studies directly probing whether the fusiform is sensitive to expression have shown mixed results ^2,13-16^. Positive findings for fusiform sensitivity to expression have led to the competing hypothesis that the division of face processing is not for identity and expression, but rather form/structure and motion ^3^. Notably though, even studies with positive results have not examined whether the same patches of the fusiform that code for identity also code for expression (see ^14^ for a study that examined both, but saw negative results for expression coding). Furthermore, some studies show the fusiform has an expression-independent identity code ^6,8^.

Beyond whether the fusiform responds differentially to expression, one key question is whether the fusiform intrinsically codes for expression or if differential responses are due to task-related and/or top-down modulation of fusiform activity ^17^. Assessing this requires a method with high temporal resolution to distinguish between early, more bottom-up biased activity, and later activity that likely involved recurrent interactions. Furthermore, a passive viewing or incidental task is required to exclude biases introduced by variable task demands across stimuli. The low temporal resolution of fMRI makes it difficult to disentangle early bottom-up processing from later top-down and recurrent processing ^8^. Some previous intracranial electroencephalography (iEEG) studies have used an explicit expression identification task, making task effects difficult to exclude ^18-20^. Those that have used an implicit task have shown mixed results regarding whether early fusiform response is sensitive to expression ^20,21^. Furthermore, iEEG studies often lack sufficient subjects and population-level analysis to allow for a generalizable interpretation.

To help mediate between these two models and clarify the role of the fusiform in facial expression perception, iEEG was recorded from 17 subjects with a total of 31 face sensitive electrodes in face sensitive patches of the fusiform gyrus while these subjects viewed faces with neutral, happy, sad, angry, and fearful expressions in a gender discrimination task. Multivariate temporal pattern analysis (MTPA) on the data from these electrodes was used to analyze the temporal dynamics of neural activity with respect to facial expression sensitivity in fusiform. In a subset of 7 subjects, identity coding was examined in the same electrodes also using MTPA. In addition to examining the overall patterns across all electrodes, the responses from and mid- and posterior fusiform, as well as the left and right hemisphere, were compared. To supplement these iEEG results, a meta-analysis of 64 neuroimaging studies was done examining facial expression sensitivity in the fusiform. The results support the view that fusiform response is sensitive to facial expression and suggest that the posterior and mid-fusiform regions play a qualitatively different role in facial expression processing.

## Results

### Electrode selection and face sensitivity

The locations of the 31 fusiform electrodes from 17 participants sensitive to faces are shown in Figure 1A and Table 1. The averaged event-related potential (ERP) and event-related broadband gamma activity (ERBB) responses (see Methods for detailed definitions of ERP and ERBB) for each category across all channels are shown in Figure 1C and Figure 1D respectively. The averaged sensitivity index (*d*’) for faces peaked at 160 ms (*d*’ = 1.22, *p* < 0.01 in every channel, Figure 1B). Consistent with previous findings ^8,22-24^, a strong sensitivity for faces was observed in fusiform around 100-400 ms after stimulus onset.

**Table 1.** MNI coordinates and facial expression sensitivity (*d’*) for all face sensitive electrodes. Electrode ID is labeled by subject number (SX) and electrode from that subject (a, b, etc.). Sensitivity to expression defined as *p* < 0.05 decoding accuracy corrected for multiple comparisons.

| Electrode ID | X (mm) | Y (mm) | Z (mm) | Peak time (ms) | Peak $d'$ | Sensitive to expressions |
| --- | --- | --- | --- | --- | --- | --- |
| S1a | 35 | -59 | -22 | 260 | 0.29 | Y |
| S1b | 33 | -53 | -22 | 150 | 0.31 | Y |
| S1c | 42 | -56 | -26 | 200 | 0.20 | N |
| S2a | 40 | -57 | -23 | 170 | 0.34 | Y |
| S3a | -33 | -44 | -31 | 580 | 0.18 | N |
| S4a | -38 | -36 | -30 | 440 | 0.12 | N |
| S5a | -38 | -36 | -20 | 300 | 0.25 | Y |
| S5b | -42 | -37 | -19 | 330 | 0.25 | Y |
| S6a | 34 | -40 | -11 | 540 | 0.24 | Y |
| S6b | 39 | -40 | -10 | 490 | 0.33 | Y |
| S7a | 36 | -57 | -21 | 100 | 0.42 | Y |
| S8a | -22 | -72 | -9 | 100 | 0.23 | Y |
| S8b | -40 | -48 | -23 | 170 | 0.38 | Y |
| S9a | 32 | -46 | -7 | 180 | 0.34 | Y |
| S9b | 36 | -48 | -8 | 160 | 0.40 | Y |
| S10a | 29 | -46 | -15 | 310 | 0.31 | Y |
| S11a | -25 | -38 | -17 | 580 | 0.36 | Y |
| S11b | -34 | -38 | -18 | 400 | 0.46 | Y |
| S11c | -49 | -37 | -20 | 430 | 0.27 | Y |
| S12a | 41 | -33 | -19 | 70 | 0.06 | N |
| S12b | 37 | -51 | -9 | 70 | 0.22 | Y |
| S12c | 35 | -59 | -4 | 80 | 0.23 | Y |
| S13a | 43 | -36 | -13 | 400 | 0.11 | N |
| S13b | 44 | -48 | -11 | 190 | 0.10 | N |
| S14a | -52 | -54 | -17 | 30 | 0.14 | N |
| S15a | -37 | -47 | -10 | 180 | 0.64 | Y |
| S16a | -39 | -45 | -11 | 160 | 0.03 | N |
| S17a | -43 | -53 | -26 | 90 | 0.20 | N |
| S17b | -46 | -50 | -28 | 110 | 0.31 | Y |
| S17c | -30 | -63 | -20 | 120 | 0.27 | Y |
| S17d | -45 | -56 | -25 | 40 | 0.13 | N |

**Figure 1.**
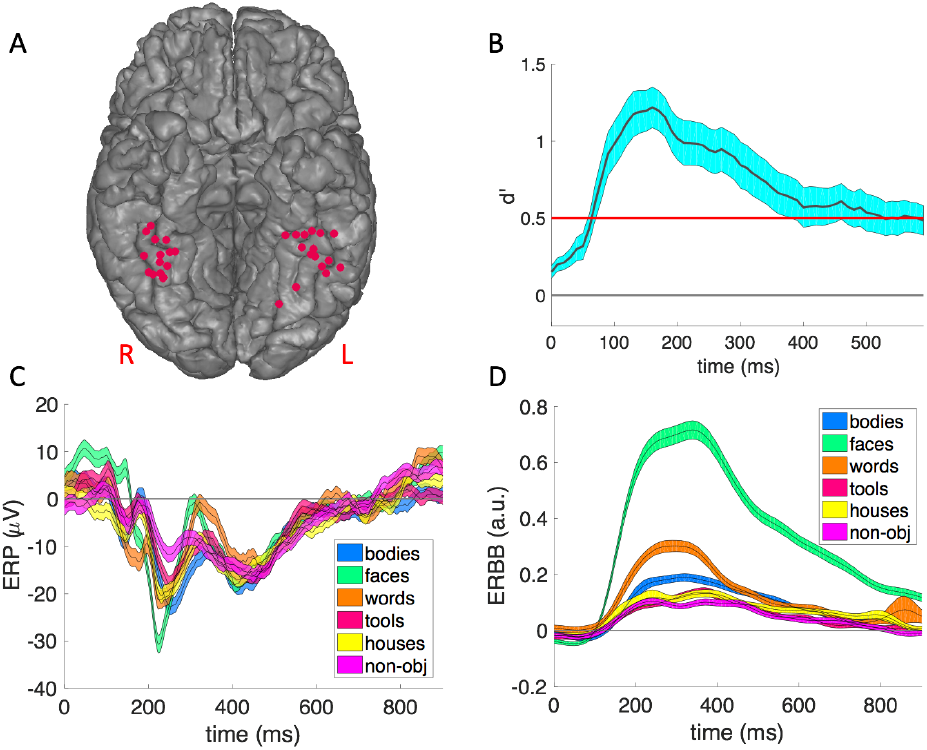
The face sensitive electrodes in the fusiform. **A)** The localization of the 31 face sensitive electrodes in (or close to) fusiform area, mapped onto a common space based on MNI coordinates. We moved depth electrode locations to the nearest location on the overlying cortical surface, in order to visualize all the electrodes. **B)** The timecourse of the sensitivity index (*d*’) for faces versus the other categories in the six-way classification averaged across all 31 fusiform electrodes. The shaded areas indicate standard error of the mean. The red line corresponds to *p* **<** 0.01 with Bonferroni correction for multiple comparisons across 60 time points. **C)** The ERP for each category averaged across all face sensitive fusiform electrodes. The shaded areas indicate standard error of the mean. **D)** The ERBB for each category averaged across all face sensitive fusiform electrodes. The shaded areas indicate standard error of the mean.

### Facial expression classification at the individual and group level

For each participant, the classification accuracy between each pair of facial expressions was estimated using 5-fold cross-validation (see Methods for details). As shown in Figure 2B, the averaged timecourse peaked at 190 ms after stimulus onset (average decoding at peak *d’* = 0.12, *p* < 0.05, Bonferroni corrected for multiple comparisons). In addition to the grand average, on the single electrode level, 21 out of the 31 electrodes from 12 out of 17 subjects showed a significant peak in their individual timecourses (*p* < 0.05, permutation test corrected for multiple comparisons). The locations of the significant electrodes are shown in Figure 2A and all electrodes are listed in Table 1.

**Figure 2.**
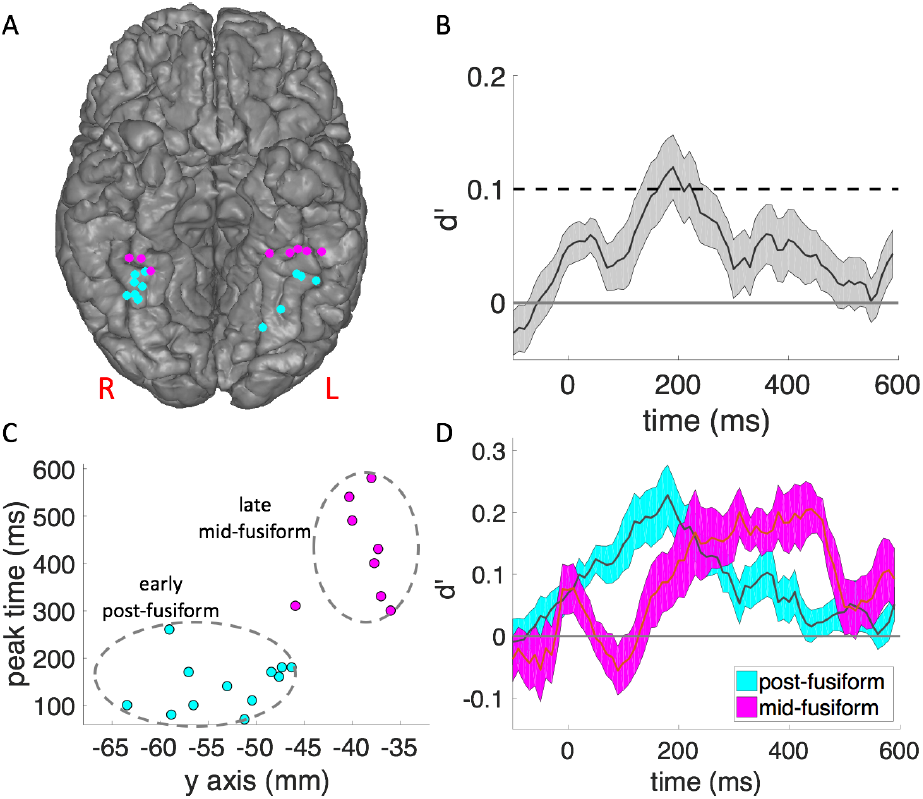
The timecourse of the facial expression classification in fusiform. **A)** The locations of the electrodes with significant face expression decoding accuracy, with the posterior fusiform group colored in cyan and the mid-fusiform group colored in magenta. **B)** The timecourse of mean and standard error for pairwise classification between different face expressions in all 31 fusiform electrodes. Dashed line: *p* = 0.05 threshold with Bonferroni correction for 60 time points [600 ms with 10 ms stepsize]). **C)** The time of the peak classification accuracy was plotted against the MNI y-coordinate for each single electrode with significant expression classification accuracy. *K*-means clustering partitions these electrodes into posterior and mid-fusiform groups. **D)** The mean and standard error for pairwise classification between different face expressions in posterior fusiform electrodes and mid-fusiform electrodes. The posterior group peaked at 180 ms after stimulus onset and the mid-fusiform group had an extended peak starting at 230 ms and extending to 450 ms (both *p* < 0.05, binomial test, Bonferroni corrected). See supplement for receiver operator characteristic (ROC) curves validating classification analysis (Figure S1).

The effect size for the mean peak expression classification is relatively low. This is in part because the electrodes consisted of two distinct populations with different timecourses (see below). Additionally, due to the variability in electrode position, iEEG effect sizes can be lower in some cases than what would be seen with electrodes optimally placed over face patches. To assess whether this was the case, we examined the correspondence between face category decoding and expression decoding based on the logic that placement closer to face patches should lead to higher face category decoding accuracy. A significant positive correlation between the decoding accuracy (*d’*) for face category and the decoding accuracy (*d’*) for facial expressions was seen (Pearson correlation *r* = 0.57, N = 21, *p* = 0.007). This suggests that electrode position relative to face patches in the fusiform can explain some of the effect size variability for expression classification. That suggests the true effect size for expression classification for optimal electrode placement may be closer to what was seen for electrodes with higher accuracy (0.4-0.6, see Table 1) rather than the mean across all electrodes.

### Spatiotemporal dynamics of facial expression decoding

The next question we addressed was whether spatiotemporal dynamics of facial expression representation in fusiform was location dependent. Specifically, we compared the dynamics of expression sensitivity between left and right hemispheres, and between posterior and mid-fusiform regions for electrodes showing significant expression sensitivity.

We first analyzed the lateralization effect for the expression coding in fusiform. The mean timecourses of decoding accuracy for left and right fusiform did not differ at the *p* < 0.05 uncorrected level at any time point (Figure S2).

In contrast substantial differences were seen in the timing and representation of expression coding between posterior and mid-fusiform. This was first illustrated by plotting the time of the peak decoding accuracy in each individual electrode against the corresponding MNI y-coordinate of the electrode (Figure 2C). A qualitative difference was seen between the peak times for electrodes posterior to approximately y = −45 compared to those anterior to that, rather than a continuous relationship between y-coordinate and peak time. This was quantified by a clustering analysis using both Bayesian information criterion (BIC) ^25^ and Silhouette analysis ^26^ (Figure S3), which both showed evidence for a cluster-structure in the data (Bayes factor > 20) with *k* = 2 as the optimal number of clusters (mean Sillhouette coefficient = 0.59). The 2 clusters corresponded to the posterior and mid-fusiform (Figure 2C; see Supplemental Information for detailed analysis on clustering and the selection of models with different values of *k*). The border between these data-driven clusters corresponds well with prior functional and anatomical evidence showing that the mid-fusiform face patch falls within a 1 cm disk centered around the anterior tip of mid-fusiform sulcus (MFS; which falls at y = −40 in MNI coordinates) with high probability ^27^. That would make the border between the mid-fusiform and posterior fusiform face patch approximately y = −45 in MNI coordinates, which is very close to the border produced by the clustering analysis (y= −45.9).

The timecourse of the posterior and mid-fusiform clusters were then examined in detail. As shown in Figure 2D, the timecourse of decoding accuracy in the posterior group peaked at 180 ms after stimulus onset and the timecourse of mid-fusiform group first peaked at 230 ms and the peak extended until approximately 450 ms after stimulus onset.

### Representational Similarity Analysis

A recent meta-analysis suggests that fusiform is particularly sensitive to the contrast between specific pairs of expressions ^28^. To examine this in iEEG data, the representation dissimilarity matrices (RDMs) for facial expressions in the early and late activity in posterior and mid-fusiform were computed (Figure 3). No contrasts between expressions showed significant differences in posterior fusiform in the late window or in mid-fusiform in the early window (*p* > 0.1 in all cases, T-test), as expected due to the corresponding low overall classification accuracy. In the early posterior fusiform, expressions of negative emotions (fearful, angry) were dissimilar to happy and neutral expressions (*p* < 0.05 in each case, T-test), but not very distinguishable from one another. In the late mid-fusiform activity, happy and neutral expressions were both distinguishable from expressions of negative emotions and from each other (*p* < 0.05 in each case, T-test). The results showed partial consistency with a previous meta-analysis based on neuroimaging studies (consistent in angry vs. neutral, fearful vs. neutral, fearful vs. happy, and fear vs sad) ^28^. However, the previous meta-analysis also reported significant contrast in fearful vs. angry and angry vs. sad, which were absent in our results.

**Figure 3.**
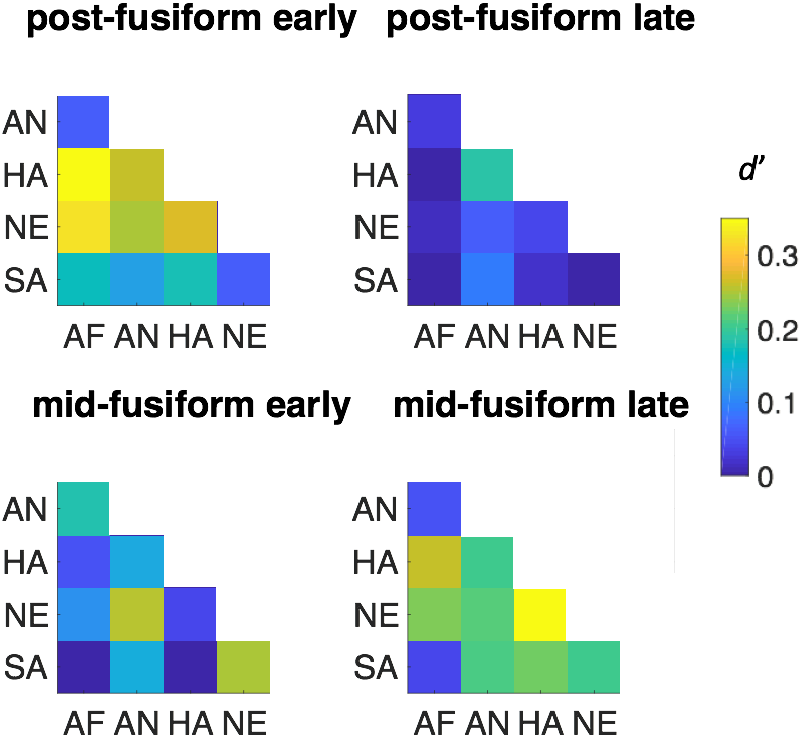
Representational similarity analysis (RSA) between the facial feature space and the representational spaces of posterior and mid-fusiform at both early and late stages. **Top row:** representational dissimilarity matrices (RDM) of posterior fusiform at early stage (left), RDM of posterior fusiform at late stage (right). **Bottom row:** RDM of mid-fusiform at early stage (left), RDM of mid-fusiform at late stage (right). Abbreviations: AF – fearful, AN – angry, HA – happy, NE – neutral, SA – sad.

One question is the degree to which the representation in fusiform reflects the physical properties of the images subjects were viewing versus a more abstract representation of emotion. To examine this question, an 17-dimensional facial feature space was constructed based on a computer vision algorithm ^29^. The features characterize structural and spatial frequency properties of each image, e.g. eye width, eyebrow length, nose height, eye-mouth width ratio, skin tone, etc. An RDM was then built between the expressions in this feature space and compared to the neural feature spaces. There was a significant correlation between posterior fusiform representation space in the early time window (Spearman’s rho = 0.24, *p* < 0.05, permutation test). The correlation between mid-fusiform representation space in the late time window and the facial feature space was smaller and did not reach statistical significance (Spearman’s rho = 0.15, *p* > 0.1, permutation test). This suggests the earlier representation reflects the physical aspects of the images more closely whereas the later representation may also reflect the emotional content more abstractly.

### Comparison to Facial Identity Classification

Given the strongly supported hypothesis the fusiform plays a central role in face identity recognition, the effect size of identity and expression coding in the fusiform was compared. Due to the relatively few repetitions of individual faces, individuation was examined in only the 7 subjects that had sufficient repetitions of each face identity allowing for multivariate classification of identity across expression; identity decoding was previously reported for 4 of these subjects in a recent study ^8^. Across the 7 total subjects (3 here and 4 reported previously), the mean peak *d’* = 0.50 for face identity decoding was significantly greater than the mean peak accuracy for facial expression decoding in the fusiform overall (t(26) = 2.64, *p* = 0.0069) as well as in both the posterior (t(18) = 2.08, *p* = 0.026) and mid-fusiform (t(13) = 2.02, *p* = 0.032) (Figure 4). With regards to the timing of identity (mean peak time = 314 ms) versus expression sensitivity, the posterior peak time for expression classification was significantly earlier than the peak time for identity (t(18) = 4.45, *p* = 0.0003), while the mid-fusiform extended peak time for expression classification overlapped with the peak time for identity.

**Figure 4.**
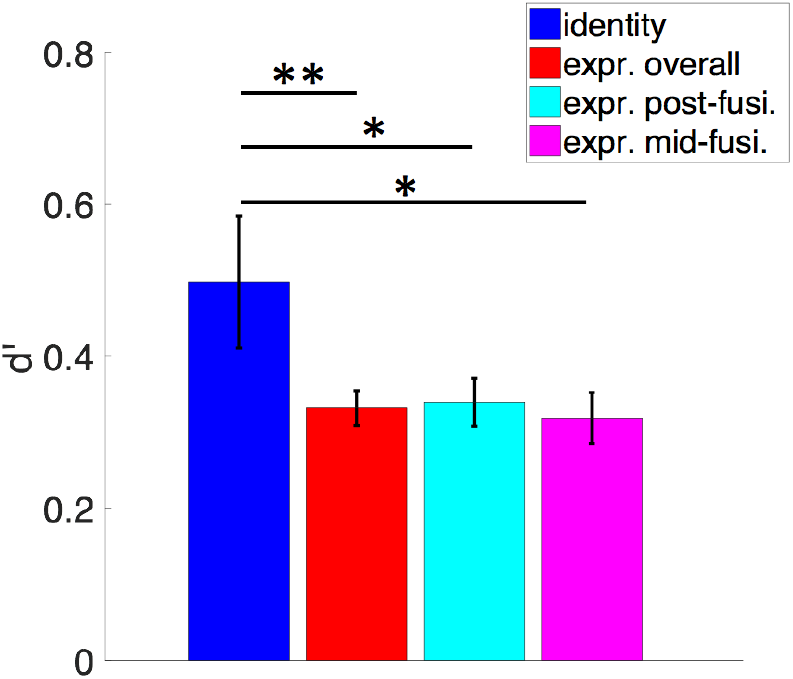
The comparison between the decoding accuracy for face identities and facial expressions. The average peak *d’* for face identity decoding (in blue) compared to the average *d’* for facial expression decoding overall (in red), in posterior fusiform (in cyan), and in anterior fusiform (in magenta). (error bar: standard error, ** *p* < 0.01, * *p* < 0.05, T-test).

## Discussion

Multivariate classification methods were used to evaluate the encoding of facial expressions recorded from electrodes placed directly in face sensitive fusiform cortex. Though the effect size for expression classification is smaller than for identity classification, the results support a role for the fusiform in the processing of facial expressions. Electrodes that were sensitive to expression were also sensitive to identity, suggesting a shared neural substrate for identity and expression coding in the fusiform. The results also show that the posterior and mid-fusiform are dynamically involved in distinct stages of facial expression processing and have different representations of expressions. The differential representation and magnitude of the temporal displacement between the sensitivity in posterior and mid-fusiform suggests these are qualitatively distinct stages of facial expression processing and not merely a consequence of transmission or information processing delay along a feedforward hierarchy.

### Fusiform is sensitive to facial expression

The results here show that the fusiform is sensitive to expression, though the effect size for classification of expression in the fusiform using iEEG is small-to-medium^1^ ^30^. The results also suggest that the same patches of the fusiform that are sensitive to expression are sensitive to identity as well. Given the variability of the effect size due to the proximity of electrode placement relative to face patches, the relative effect size may be more informative than the absolute effect size. The magnitude for facial expression classification is approximately half what was seen for face identity classification. This suggests that while fusiform contributes to facial expression perception, it is to a lesser degree than face identity processing. Greater involvement in identity than expression perception is expected for a region involved in structural processing of faces because identity relies on this information more than expression as expression perception also relies on facial dynamics. These results support models that hypothesize fusiform involvement in form/structural processing, at least for posterior fusiform (see discussion on spatially and temporally segregated stages of processing below), which can support facial expression processing ^3,5^. These results do not support models that hypothesize a strong division between facial identity and expression processing ^1,2^.

To test what a brain region codes for one must examine its response for early, bottom-up activation during an incidental task or passive viewing ^17^, otherwise it is difficult to disentangle effects of task demands and top-down modulation. Indeed, previous studies have demonstrated that extended fusiform activity, particularly in the broadband gamma range, is modulated by task-related information ^8,31^. Some previous iEEG studies of expression coding in the fusiform have used an explicit expression judgment task and examined only broadband gamma activity, making it difficult to draw definitive conclusions about fusiform expression coding from these results ^18,19^. One previous study that used an implicit task did not show evidence of expression sensitivity during the early stage of activity in the fusiform ^20^; another did show evidence of expression sensitivity, though it reported results only from a single subject ^21^. The results here show in a large iEEG sample that the early response of the fusiform most sensitive to bottom-up processing is modulated by expression, at least for the posterior fusiform.

The effect size for facial expression classification is consistent with mixed findings in the neuroimaging literature for expression sensitivity in the fusiform ^2,13-16,19^. IEEG generally has greater sensitivity and lower noise than non-invasive measures of brain activity. Methods with lower sensitivity, such as fMRI, would be expected to have a substantial false negative rate for facial expression coding in the fusiform. To quantify fMRI sensitivity to expression we performed a meta-analysis on 64 studies. Of these studies, 24 reported at least one expression sensitive loci in the fusiform. However, at the meta-analytic level, no significant cluster of expression sensitivity was seen in the fusiform after whole brain analysis (see Supplemental Table S1, S2, and Figure S4). Thus, consistent with the iEEG effect size for expression decoding in the fusiform seen here, there is some suggestion in the fMRI literature for expression sensitivity in the fusiform, but it is small and does not achieve statistical significance at the whole brain level.

### Multiple, spatially and temporally segregated stages of face expression processing in the fusiform

Using a data-driven analysis, posterior and mid-fusiform face patches were shown to contribute differentially to expression processing. The dividing point between post-fusiform and mid-fusiform electrodes found in a data-driven manner is consistent with the anatomical border for the posterior and mid-fusiform face patches previously described, suggesting a strong coupling between the anatomical and functional divisions in fusiform ^27^. While posterior and mid-fusiform have been shown to be cytoarchitectonically distinct regions each with separate face sensitive patches ^27,32,33^, functional differences between these patches have remained elusive in the literature. The results here suggest that these anatomical and physiological distinctions correspond to functional distinctions in the role of these areas in face processing, as reflected in qualitatively different temporal dynamics in these regions for facial expression processing. Specifically, posterior fusiform participates in a relatively early stage of facial expression processing that may be related to structural encoding of faces while mid-fusiform demonstrates a distinct pattern of extended dynamics and participates in a later stage of processing that may be related to a more abstract and/or multifaceted representation of expression and emotion. These results support the revised model of fusiform function that posits the fusiform contributes to structural encoding of facial expression during the initial stages of processing ^3,5^, with the notable addition that it may be that only posterior fusiform contributes to this processing.

The early time period of expression sensitivity in posterior fusiform overlaps with strong face sensitive activity measured non-invasively around 170 ms after viewing a face, which is thought to reflect structural encoding of face information ^22,23,34-36^. Face sensitive activity in this time window has been shown to be insensitive to attention and is thought to reflect a “rapid, feed-forward phase of face-selective processing.”^37^ Additionally, a face adaptation study showed that activity in this window reflects the actual facial expression rather than the perceived (adapted) expression ^38^. Consistent with these previous findings, the RSA results here show that the early posterior activity is more closely correlated to the physical features of the face that relate to facial expression compared to later, mid-fusiform activity.

The expression sensitivity in mid-fusiform onset began later than the posterior fusiform (around 230 ms), and remained active until ~450 ms after viewing a face. Face sensitive activity in this time window has been shown to be sensitive to face familiarity and to attention ^39,40^. Previous studies and the results presented here show that face identity can be decoded from the activity in this later time window in mid-fusiform ^8,41^ and reflects a distributed code for identity among regions of the face processing network ^42^. Additionally, the previously mentioned face adaptation study showed that activity in this window reflects the subjectively perceived facial expression after adaptation ^38^. The RSA analysis here showed that the activity in this time window in mid-fusiform was not significantly correlated with physical features of the face and therefore may reflect more subjective expression perception. Taken together, these results suggest the mid-fusiform expression sensitivity in this later window reflect a more abstract and subjective representation of expression and may be related to integration of multiple face cues, including identity and expression. This abstract and multifaceted representation is likely to reflect processes arising from interactions across the face processing network ^4^.

To conclude, the results presented here support the hypothesis that the fusiform contributes to expression processing ^3,5^. The finding that the same part of the fusiform is sensitive to both identity and expression contradicts models that hypothesize separate pathways for their processing ^1,2^ and instead supports the hypothesis that form and motion are the critical functional separation ^3^. The results also show there is a qualitative distinction between face processing in posterior and mid-fusiform, with each contributing to temporally and functionally distinct stages of expression processing. This distinct contribution of these two fusiform patches suggest that the structural and cytoarchitectonic differences between posterior and mid-fusiform are associated with functional differences between the contributions of these areas to face perception. Taken together, the results here illustrate the dynamic role the fusiform plays in multiple stages of facial expression processing.

## Methods

### Participants

The experimental protocols were approved by the Institutional Review Board of the University of Pittsburgh. Written informed consent was obtained from all participants.

17 human subjects (8 male, 9 female) underwent surgical placement of subdural electrocorticographic electrodes or stereoelectroencephalography (together electrocorticography and stereoelectroencephalography are referred to here as iEEG) as standard of care for seizure onset zone localization. The ages of the subjects ranged from 19 to 65 years old (mean = 37.9, SD = 12.7). None of the participants showed evidence of epileptic activity on the fusiform electrodes used in this study nor any ictal events during experimental sessions.

### Experiment design

In this study, each subject participated in two experiments. Experiment 1 was a functional localizer experiment and Experiment 2 was a face perception experiment. The experimental paradigms and the data pre-processing methods were similar to those described previously by Ghuman and colleagues ^8^.

#### Stimuli

In Experiment 1, 180 images of faces (50% male), bodies (50% male), words, hammers, houses, and phase scrambled faces were used as visual stimuli. Each of the six categories contained 30 images. Phase scrambled faces were created in MATLAB^TM^ by taking the 2-dimensional spatial Fourier spectrum of each of the face images, extracting the phase, adding random phases, recombining the phase and amplitude, and taking the inverse 2-dimensional spatial Fourier spectrum.

In Experiment 2, face stimuli were taken from the Karolinska Directed Emotional Faces stimulus set. Frontal views and 5 different facial expressions (fearful, angry, happy, sad, and neutral) from 70 faces (50% male) in the database were used, which yielded a total of 200 unique images.

#### Paradigms

In Experiment 1, each image was presented for 900 ms with 900 ms inter-trial interval during which a fixation cross was presented at the center of the screen (~ 10° × 10° of visual angle). At random, 1/3 of the time an image would be repeated, which yielded 480 independent trials in each session. Participants were instructed to press a button on a button box when an image was repeated (1-back).

In Experiment 2, each face image was presented for 1500 ms with 500 ms inter-trial interval during which a fixation cross was presented at the center of the screen. This yielded 200 independent trials per session. Faces subtended approximately 5 degrees of visual angle in width. Subjects were instructed to report whether the face was male or female via button press on a button box.

Paradigms were programmed in MATLAB^TM^ using Psychtoolbox and custom written code. All stimuli were presented on an LCD computer screen placed approximately 150 cm from participants’ heads.

All of the participants performed one session of Experiment 1. 9 of the subjects performed one session of Experiment 2, and the other 8 participants performed two or more sessions of Experiment 2.

### Data analysis

#### Data preprocessing

The electrophysiological activity was recorded using iEEG electrodes at 1000 Hz. Single-trial potential signal was extracted by band-passing filtering the raw data between 0.2-115 Hz using a forth order Butterworth filter to remove slow and linear drift, and high frequency noise. The 60 Hz line noise was removed using a forth order Butterworth filter with 55-65 Hz stop-band. Power spectrum density (PSD) at 2 – 100 Hz with bin size of 2 Hz and time-step size of 10 ms was estimated for each trial using multi-taper power spectrum analysis with Hann tapers, using FieldTrip toolbox ^43^. For each channel, the neural activity between 50 and 300 ms prior to stimulus onset was used as baseline, and the PSD at each frequency was then z-scored with respect to the mean and variance of the baseline activity to correct for the power scaling over frequency at each channel. The broadband gamma signal was extracted as mean z-scored PSD across 40-100 Hz. Event-related potential (ERP) and event-related broadband gamma signal (ERBB), both time-locked to the onset of stimulus from each trial, were used in the following data analysis.

To reduce potential artifacts in the data, raw data were inspected for ictal events, and none were found during experimental recordings. Trials with maximum amplitude 5 standard deviations above the mean across all the trials were eliminated. In addition, trials with a change of more than 25 μV between consecutive sampling points were eliminated. These criteria resulted in the elimination of less than 1% of trials.

#### Electrode localization

Coregistration of grid electrodes and electrode strips was adapted from the method of Hermes, et al. ^44^. Electrode contacts were segmented from high resolution post-operative CT scans of patients coregistered with anatomical MRI scans before neurosurgery and electrode implantation. The Hermes method accounts for shifts in electrode location due to the deformation of the cortex by utilizing reconstructions of the cortical surface with FreeSurfer^TM^ software and co-registering these reconstructions with a high-resolution post-operative CT scan. SEEG electrodes were localized with Brainstorm software ^45^ using post-operative MRI co-registered with pre-operative MRI images.

#### Electrode selection

Face sensitive electrodes were selected based on both anatomical and functional constraints. Anatomical constraint was based upon the localization of the electrodes on the reconstruction using post-implantation MRI. In addition, multivariate temporal pattern analysis (MTPA) was used to functionally select the electrodes that showed sensitivity to faces, comparing to other conditions in the localizer experiment (see below for MTPA details). Specifically, three criterions were used to screen and select the electrodes of interest: (1) electrodes of interest were restricted to those that were located in or near the fusiform gyrus; (2) electrodes were selected such that their peak 6-way classification *d’* score for faces (see below for how this was calculated) exceeded 0.5 (*p* < 0.01 based on a permutation test, as described below); (3) electrodes were selected such that the peak amplitude of the mean event related potential (ERP) and/or mean event related broadband gamma signal (ERBB) for faces was larger than the peak of mean ERP and/or ERBB for the other non-face object categories in the time window of 0 – 500 ms after stimulus onset.

#### Multivariate temporal pattern analysis (MTPA)

Multivariate methods were used instead of traditional univariate statistics because of their superior sensitivity ^8,46-48^. In this study, MTPA was applied to decode the coding of stimulus condition in the recorded neural activity. The timecourse of the decoding accuracy was estimated by classification using a sliding time window of 100 ms. Both ERP and ERBB signals in the time window are combined as input features for the MTPA classifier, which yields 110 temporal features in each case (100 voltage potentials for ERP and 10 normalized mean power-spectrum density for ERBB). The 110 dimensional data were then used as input for the classifier. The goal of the classifier was to learn the patterns of the data distributions in such 110-dimensional space for different conditions and to decode the conditions of the corresponding stimuli from the testing trials. The classifier was trained on each electrode of each subject separately to assess the electrode sensitivity to faces and facial expressions. For Experiment 1, it was a 6-way classification problem and we specifically focused on the sensitivity of face category against other non-face categories. Therefore, we used the sensitivity index (*d*’) for face category against all other non-face category as the metric of face sensitivity. *d*’ was calculated as Z(true positive rate) – Z(false positive rate) where Z is the inverse of the Gaussian cumulative distribution function. *d*’ was used because it is an unbiased measure of effect size and one that takes into both the true positive and false positive rates. It also has the advantage that it is an effect size measure that has similar interpretation as Cohen’s *d* ^30,49^ while also being applicable to multivariate classification. In addition, we provide full receiver-operator characteristic (ROC) curves for completeness and as validation of *d*’ values. For Experiment 2, averaged pair-wise classification between every possible pair of facial expressions (10 pairs in total) was used.

The choice of the classifier is an empirical problem. The performance of the classifier depends on whether the assumptions of the classifier approximate the underlying truth of the data. Additionally, the complexity of the model and the size of the dataset affect performance (bias-variance trade-off). In this study, we employed Naïve Bayes (NB) classifiers, which assumes that each of the input features are conditionally independent from one another, and are Gaussian distributed. The classification accuracy of the classifier was estimated through 5-fold cross-validation. Specifically, all the trials were randomly and evenly spited into five folds. In each cross-validation loop, the classifier was trained based on four folds and the performance was evaluated on the left out fold. The overall performance was estimated by averaging cross all the 5 cross-validation loops. In general, different classifiers gave similar results. Specifically, we evaluated the performance of different classifiers (NB, support vector machines, and random forests) on a small subset of the data, and NB classifier tended to perform better than other commonly used classifiers in the current experiment, but other classifiers also gave similar results. In addition, our previous experience ^48^ with similar datasets also suggested that NB performed reasonably well in such classification analysis. We therefore used NB throughout the work presented here. The advantage of the Naïve Bayes classifier in the current study is likely due to intrinsic properties of the high dimensional problem ^50^ that make a high-bias low-variance classifier (i.e. NB classifier) preferable compared to the low-bias high-variance classifiers (i.e. support vector machines).

#### Permutation testing

Permutation testing was used to determine the significance of the sensitivity index *d’*. For each permutation, the condition labels of all the trials were randomly permuted and the same procedure as described above was used to calculate the *d*’ for each permutation. The permutation was repeated for a total of 1000 times. The *d*’ of each permutation was used as the test statistic and the null distribution of the test statistic was estimated using the histogram of the permutation test.

#### *K*-means clustering

*K*-means clustering was used to cluster the electrodes into groups based on both functional and anatomical features ^26^. Specifically, we applied *k*-means clustering algorithm to the electrodes in a 2D feature space of MNI y-coordinate and the peak classification accuracy time. Note that each dimension was normalized through z-scoring in order to account for different scales in space and time. See Supplemental Information for detailed analysis using Bayesian information criterion and Silhouette analysis for model selection.

#### Facial feature analysis

The facial features from the stimulus images were extracted following the similar process as ^8^. Anatomical landmarks for each picture were first determined by IntraFace ^29^, which marks 49 points on the face along the eyebrows, down the bridge of the nose, along the base of the nose, and outlining the eyes and mouth. Based on these landmarks we calculated 17 facial feature dimensions listed in Table S3. The values for these 17 feature dimensions were normalized by subtracting the mean and dividing by the standard deviation across the all the pictures. The mean representation of each expression in facial feature space was computed by averaging across all 70 faces of the same expression.

#### Representational similarity analysis (RSA)

RSA was used to analyze the neural representational space for expressions ^51^. With pair-wise classification accuracy between each pair of facial expressions, we constructed the representational dissimilarity matrix (RDM) of the neural representation of facial expressions, with the element in the i-th column of the j-th row in the matrix corresponding to the pairwise classification accuracy between the i-th and j-th facial expressions. The corresponding RDM in the facial feature space was constructed by assessing the Euclidean distance between the i-th and j-th facial expressions in the 17-dimensional facial feature space.

## Acknowledgements

We thank the patients for participating in the iEEG experiments and the UPMC Presbyterian epilepsy monitoring unit staff and administration for their assistance and cooperation with our research. We thank Michael Ward, Ellyanna Kessler, Vincent DeStefino, Shawn Walls, Roma Konecky, Nicolas Brunet, and Witold Lipski for assistance with data collection and thank Ari Kappel and Matthew Boring for assistance with electrode localization. This work was supported by the National Institute on Drug Abuse under award NIH R90DA023420 (to YL), the National Institute of Mental Health under award NIH R01MH107797 and NIH R21MH103592 (to ASG), and the National Science Foundation under award 1734907 (to ASG). The content is solely the responsibility of the authors and does not necessarily represent the official views of the National Institutes of Health or the National Science Foundation.

## Supplement Information

### Methods

#### Clustering analysis

We applied *k*-means clustering to the electrodes in the 2D space of MNI y-coordinate and peak classification time for facial expressions, with different values of *k*, and evaluated the model performance by computing the Bayes information criterion (BIC) and the mean Silhouette coefficient (SC) across all points.

Following ^1,2^, the BIC was estimated using Schwartz criterion. Specifically, 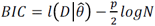, where 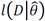 is the log-likelihood of the data under the assumption of *k*-means (spherical Gaussian) taken at the maximum likelihood estimation of parameters 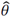, *p* is the total number of parameters in the model, and *N* is the total number of data points.

Following ^3^, the Silhouette value for the i-th point was computed as *S*_*i*_ =(*b*_*i*_ – *a*_*i*_)/max(*a*_i_, *b*_*i*_), where *a*_i_ is the average within cluster distance for the i-th point, and *b*_i_ is the minimum average between cluster distance for the i-th point (minimized over all other clusters). The mean SC was then estimated by averaging the Silhouette value over all data points.

#### Meta-analysis

Activation likelihood estimation (ALE, ^4,5^) was used for the meta-analysis of the neuroimaging literature. We first searched the online database of neuroimaging studies on Neurosynth.org and found around 300 imaging studies with the keyword “facial expressions”. We then further narrowed the list down to 64 fMRI by only including the studies that had a direct full brain mapping by contrasting between emotional facial expressions, e.g. fear vs neutral, happy vs sad, etc. We only took into account the reported activation foci for the contrast between facial expressions. Then all of the activation foci in those relevant full brain map results were collected and extracted as 3D coordinates in MNI space. In the ALE, each of the extracted foci was assigned as the center of a Gaussian distribution, whose variance was scaled by the number of subjects in the corresponding experiment. These Gaussian distributions were then combined to build a full brain map of ALE. The ALE map was corrected for multiple comparison using cluster-based permutation test. Then we performed a spatial permutation test with 1000 permutations to construct a null distribution of the full brain activation. The ALE and the corresponding statistical analysis were performed based on GingerALE 2.3.6 ^6,7^.

## Results

### Selection of models for *k-*means clustering

We applied *k*-means clustering to the electrodes in the 2D space of MNI y-coordinate and peak classification time for facial expressions, with different values of *k*, and evaluate the model performance by computing the Bayes information criterion (BIC) and the mean Silhouette coefficient (SC) across all points.

As shown in Figure S3, for *k* = 1, BIC = −61.28; for *k* = 2, BIC = −54.63.Therefore, Bayes factor between the hypothesis (H0) that there is a cluster structure (*k* =2) and the null hypothesis (H0) that there is no cluster structure (*k* = 1) can be approximated as 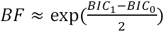. This approximation yields a *BF* > 20, which suggests a strong evidence of H1 over H0. In other words, there is a strong clustering structure in the data.

Moreover, for *k* = 2, BIC = −54.63, the mean SC = 0.601; for *k* = 3, BIC = −56.29, the mean SC = 0.490; for *k* = 4, BIC = −58.54, mean SC = 0.428. Both BIC and mean SC suggest that *k* = 2 is the optimal number of clusters. Therefore, *k* = 2 was used in the study.

### Meta-analysis of the neuroimaging literature

In the broad neuroimaging literature, we found 64 fMRI studies with full brain contrasts between face expressions (See Table S1). Among the 64 studies, 24 studies report at least one significant focus of fusiform sensitivity to differences in expressions (See Figure S4 for activation map). A total of 999 significant foci were reported in those experiments for contrasts between different facial expressions (Figure S4). A full brain activation likelihood estimation (ALE) was performed and significance was assessed using a cluster-based permutation test. 4 significant clusters were found at the *p* < 0.01 threshold, none of which included the fusiform. The MNI coordinates for the center and the corresponding label names of the 4 clusters are shown in Table S2.

---

**Figure S1.**
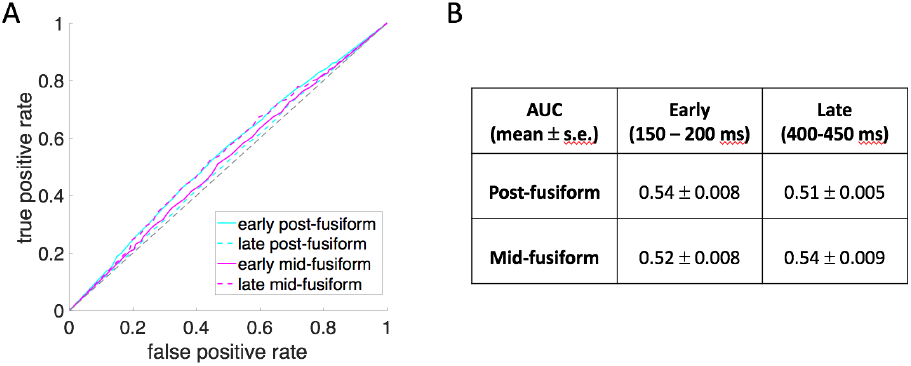
The mean ROC curve and area-under-curve (AUC) for posterior fusiform electrodes and mid-fusiform electrodes at early (150-200 ms after stim onset) and late stage (400-450 ms after stim onset).

**Figure S2.**
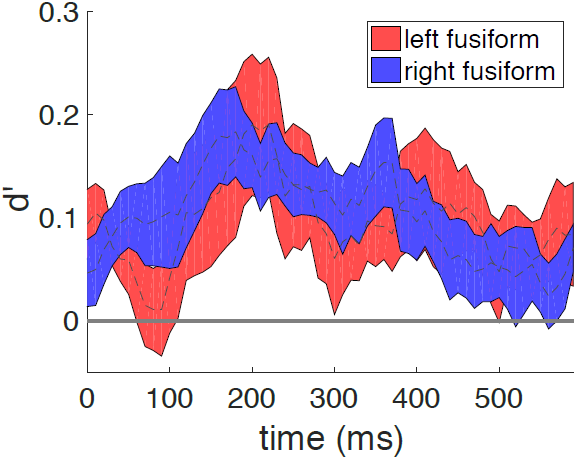
The mean and standard error for classification between different face expressions in left and right fusiform electrodes. The timecourse of the left fusiform peaked at 220 ms after stimulus onset with mean *d’* = 0.19, and the timecourse of the right fusiform peaked at 180 ms after stimulus onset with mean *d’* = 0.18 (both *p* < 0.05, binomial test, Bonferroni corrected).

**Figure S3.**
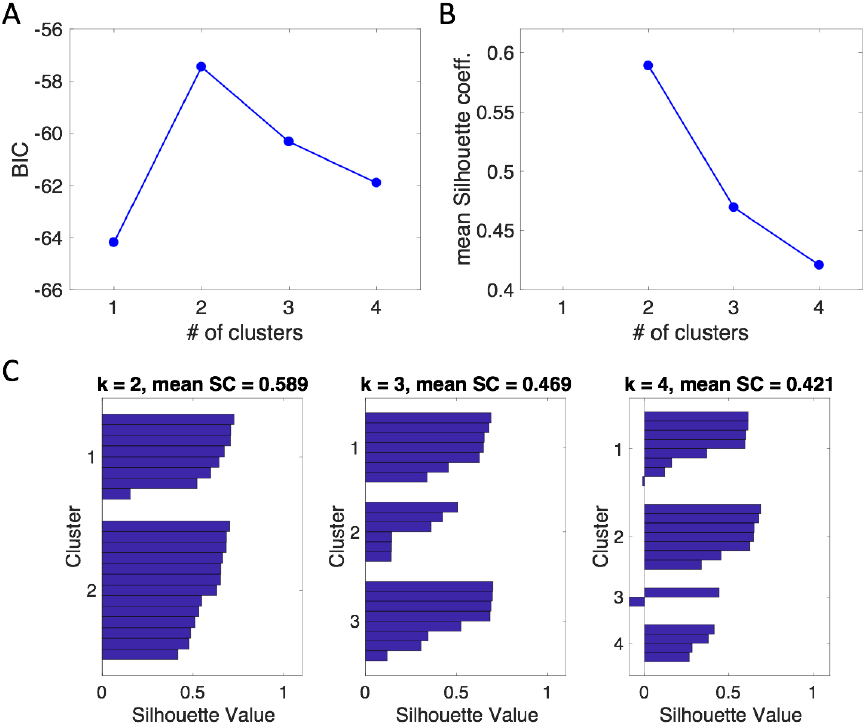
Clustering analysis. **A)** BIC of *k*-means models with different values of *k* (*k* = 1, 2, 3, 4). **B)** Mean SC of *k*-means models with different values of *k* (*k* = 2, 3, 4, note that SC is not applicable for *k* = 1). **C)** The distribution of Silhouette Coefficients (SC) with different values of *k* in *k*-means clustering. From left to right, *k* = 2, 3, and 4.

**Figure S4.**
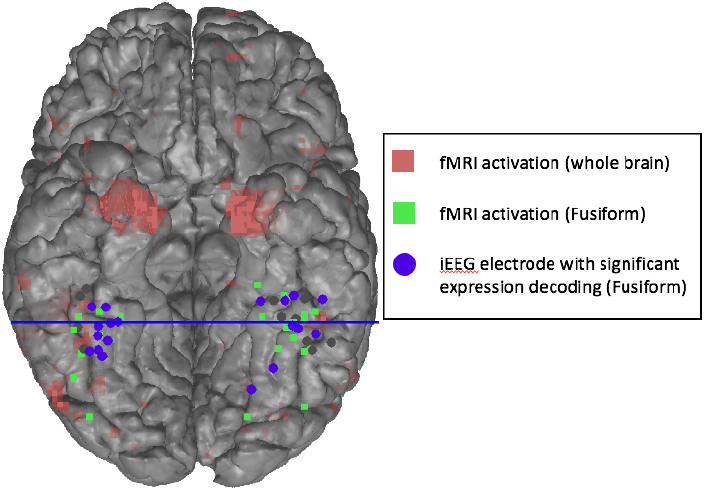
Activation map for facial expressions. (Red) Whole brain activation map from all 64 relevant fMRI studies. (Green square) Voxels in fusiform reported in 24/64 of the fMRI studies that have significant contrast between facial expressions. (Blue dots) iEEG electrodes in fusiform that have significant facial expression decoding. (Blue line) the border between posterior and mid-fusiform clusters based upon clustering analysis in the iEEG electrodes

**Table S1.**
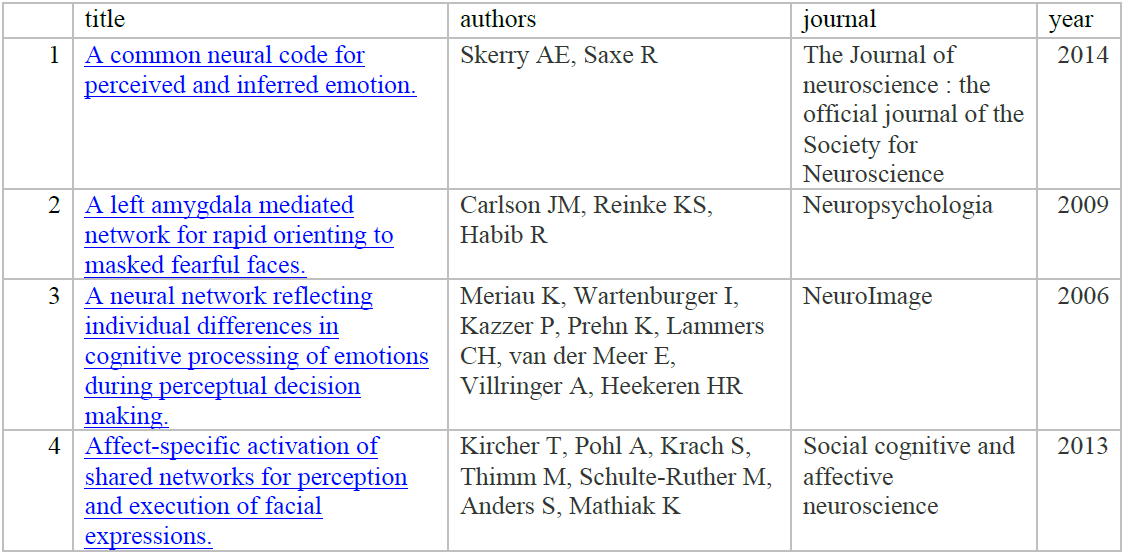
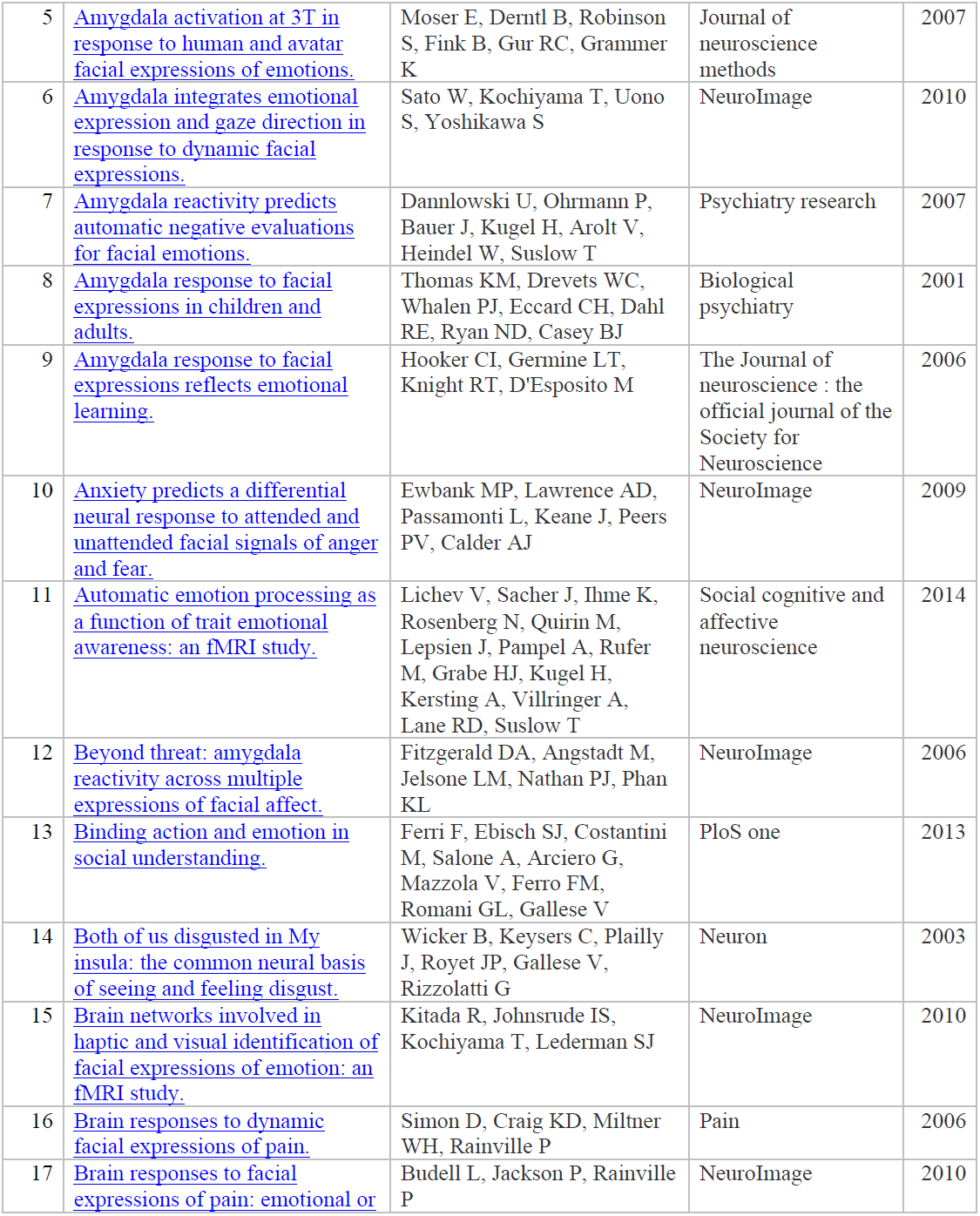
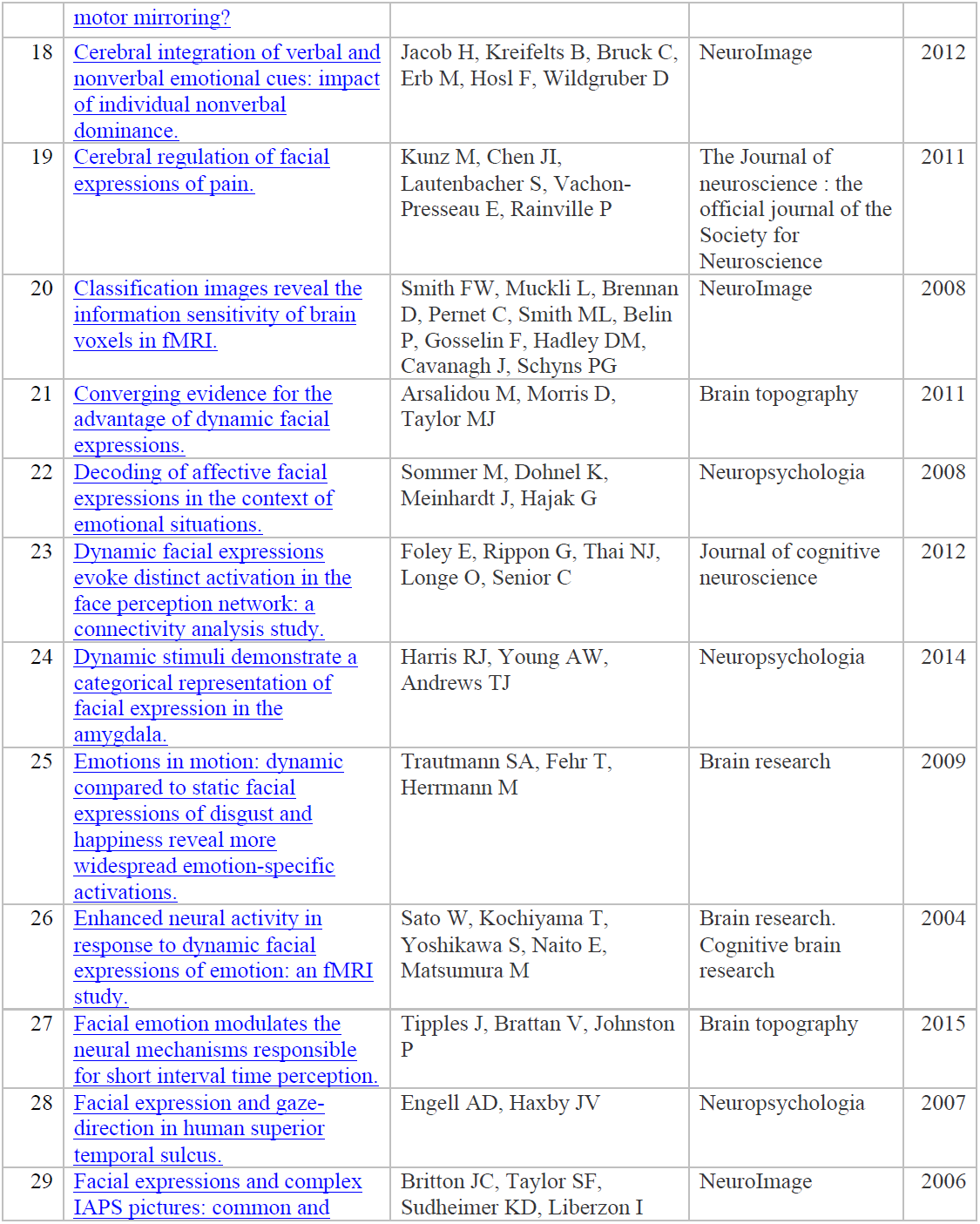
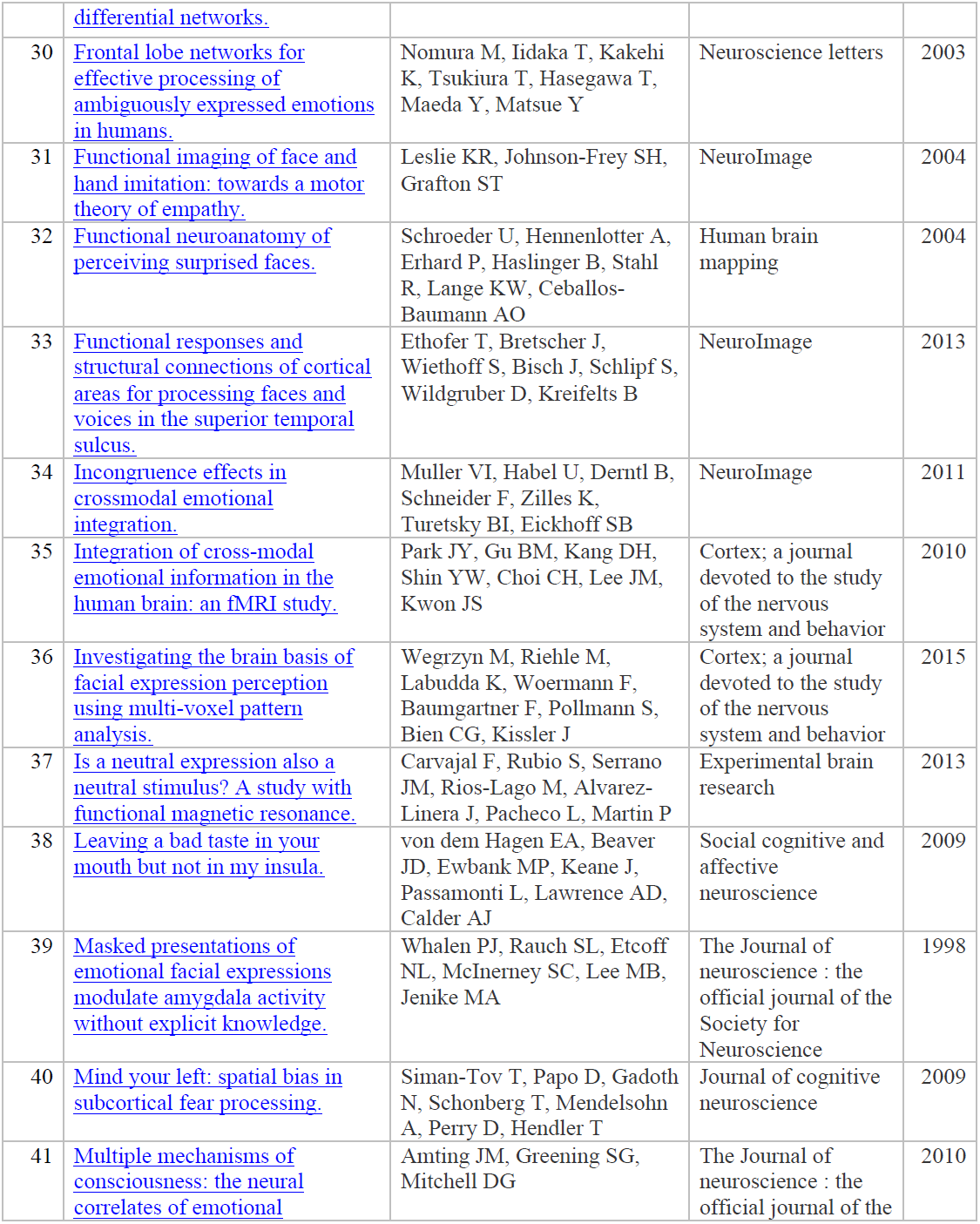
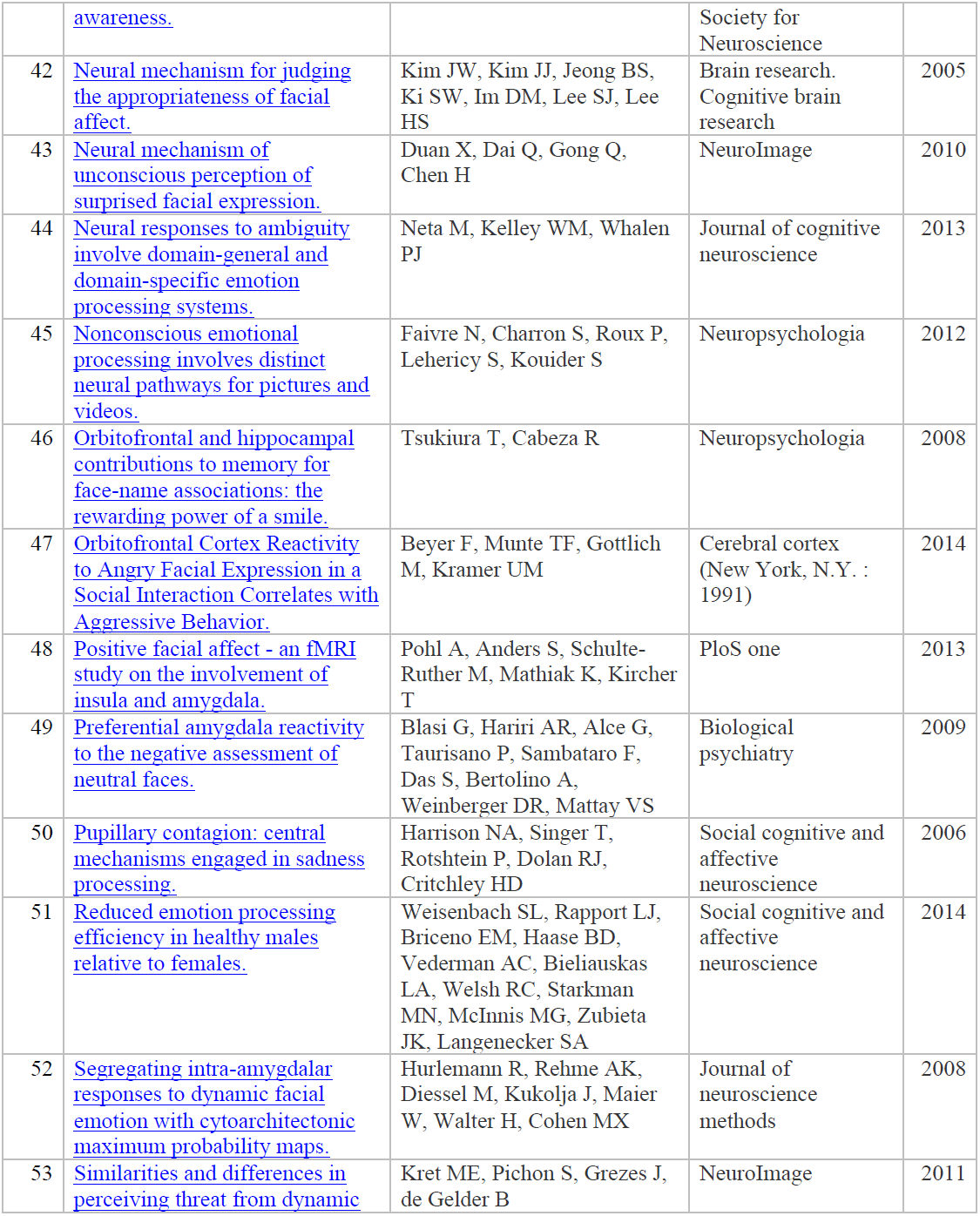
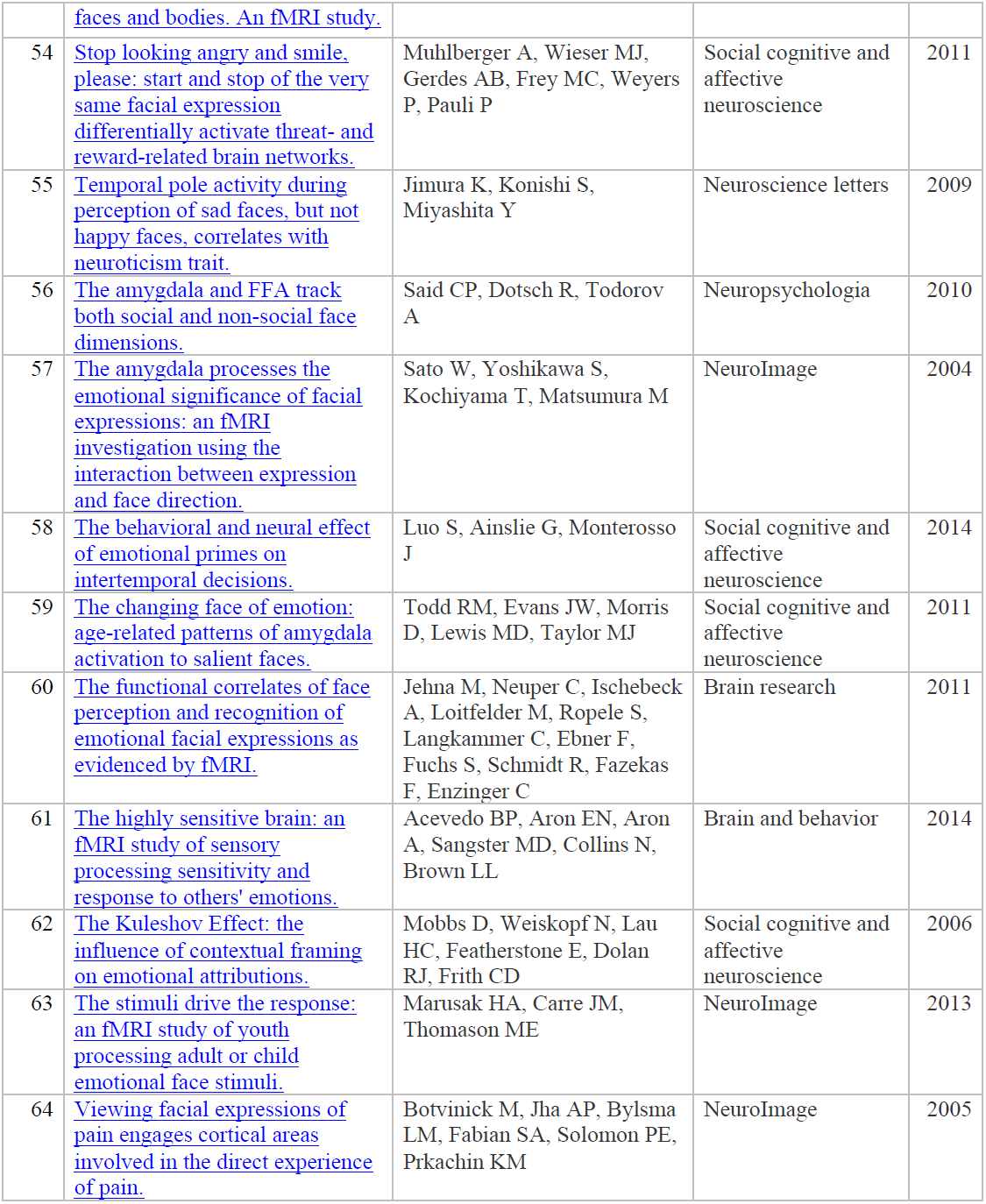
A summary list for the 64 neuroimaging studies included in the meta-analysis

**Table S2.** the MNI coordinates for the weighted center, volume, and the corresponding label name of the significant clusters in the ALE map from meta-analysis

| Cluster # | X | Y | Z | Volume (mm<sup>3</sup>) | Lateralization | Label |
| --- | --- | --- | --- | --- | --- | --- |
| 1 | 23 | -3 | -18 | 6088 | right | amygdala |
| 2 | -22 | -4 | -18 | 4640 | left | amygdala |
| 3 | 56 | -42 | 5 | 2520 | right | middle/superior temporal gyrus |
| 4 | 3 | 11 | 53 | 1464 | right | superior frontal gyrus |

**Table S3.** 17 features used for the facial feature space

| Feature # | Feature name |
| --- | --- |
| 1 | eyebrow length |
| 2 | inter-eyebrow distance |
| 3 | eye width |
| 4 | inter-eyes distance |
| 5 | vertical distance between eyes and nosetip |
| 6 | horizontal length of the nose |
| 7 | distance between nose and upper lip |
| 8 | face height |
| 9 | face width |
| 10 | eye height |
| 11 | width of the mouth |
| 12 | intense of red on cheeks |
| 13 | intense of green on cheeks |
| 14 | intense of blue on cheeks |
| 15 | contrast polarity between eyes and nose |
| 16 | eye area |
| 17 | eye mouth ratio |

## Footnotes

1 *d’* is on the same scale as Cohen’s d and they are equivalent when the data is univariate Gaussian, so *d’*s between 0.2-0.5 are “small” and 0.5-0.8 are medium.

